# Information Content of Trees: Three-taxon Statements Inference Rules and Dependency

**DOI:** 10.1101/2020.06.08.141515

**Authors:** Valentin Rineau, René Zaragüeta, Jérémie Bardin

## Abstract

The three-taxon statement is the fundamental unit of rooted trees in Cladistics, stating that for three terminal taxa, two are more related to each other than to a third. Because of their fundamental role in phylogenetics, three-taxon statements are present in methodological research of various disciplines in evolutionary biology, as in consensus methods, supertree methods, species-tree methods, distance metrics, and even phylogenetic reconstruction. However, three-taxon statements methods are subject to important flaws related to information redundancy. We aim to study the behavior of three-taxon statements and the interactions among them in order to enhance their performance in evolutionary studies. We show here how specific interactions between three-taxon statements are responsible of the emergence of redundancy and dependency within trees, and how they can be used for the improvement of weighting procedures. Our proposal is subsequently empirically tested in the supertree framework using simulations. We show that three-taxon statements using fractional weights perform drastically better than classical methods such as MRP or methods using unweighted statements. Our study shows that appropriate fractional weighting of three taxon statements is of critical importance for removing redundancy in any method using three-taxon statements, as in consensus, supertrees, distance metrics, and phylogenetic or biogeographic analyses.

---

Phylogenetic trees are at the heart of most evolutionary biology studies, and their reliability is of critical importance. It has long been recognised in the phylogenetic setting the need to understand and manipulate the information contained in phylogenies. Any tree (e.g. taxon tree, gene tree, biogeographic tree) can be fruitfully decomposed into subunits of several kinds (Adams, 1986; Vach, 1994), as components (rooted subtrees with all taxa and a single informative node; Nelson & Platnick, 1981, Wilkinson et al., 2004), three-taxon statements (3ts thereafter; rooted subtrees with three taxa and a single informative node; Adams, 1986), splits (bipartitions of a set of taxa; Farris, 1970), or quartets (bipartitions of 4 taxa; Strimmer and von Haeseler, 1996). These subunits are considered in the phylogenetic framework by several authors as information bricks (e.g. Mickevich & Platnick, 1989; Nelson, 1979; Nelson & Ladiges, 1992; Wilkinson, 1994a), and can be used to measure the information content of a phylogenetic tree.

Here we focus on rooted trees. Within a rooted tree, the 3ts (see Nelson & Ladiges, 1991a, 1991b; Nelson & Platnick, 1991; Rineau and Prin, 2021) take a special place as they represent the minimal (rooted) phylogenetic information. A 3ts is a statement in which given three distinct taxa, two are related compared to a third one. For example, given three taxa *a, b* and *c, a* and *b* are more closely related to each other than either is to *c*: *c*(*ab*). These particular types of relationships can also be found under other names in the phylogenetic literature, such as “triad” (Adams, 1986: 302), “triplet” (Wilkinson, 1994a), or still “rooted triple” (Bryant, 2003) in the consensus and supertree literature (note that we avoid here these denominations that may potentially confuse two distinct mathematical objects that are ternary relations of degree of equivalence one the one hand and sets of three elements on the other hand).

Because of their atomic nature, 3ts are at the heart of many methods. For example, the method of phylogenetic reconstruction commonly referred to as three-taxon analysis (Nelson and Ladiges, 1991b; Nelson and Platnick, 1991) decomposes hierarchical characters into 3ts matrices and then selects the cladograms that are congruent with the maximal amount of those 3ts (Zaragüeta et al., 2012). The same principle is applied in cladistic biogeography and the construction of areagrams (i.e. cladograms of biogeographic areas; Nelson and Ladiges, 1991a). The use of 3ts in the supertree framework was first proposed by Williams (2004).

Currently, several supertree methods (Ranwez et al., 2010; Sevillya et al., 2016; Dannenberg et al., 2019) explicitly handle 3ts. Another use of 3ts is also for consensus tree representations (Aho et al., 1981; Adams, 1986; Nelson & Ladiges, 1994; Wilkinson, 1994a; Bryant, 2003; Cao et al., 2009). The 3ts permit to precisely quantify the information content of a phylogenetic structure (Mickevich and Platnick, 1989; Nelson and Ladiges, 1992; Williams and Humphries, 2003). The retention index based on 3ts (Kitching et al., 1998) allows measuring the proportion of hierarchical relationships that are found in the optimal cladograms. 3ts-based distance metrics have been used to compare topologies among sets of phylogenetic trees (Grand et al., 2013; Kuhner and Yamato, 2015). Finally, 3ts are also explicitly used for coalescent-based species tree estimation from gene tree (Liu et al., 2010; Islam et al., 2020), and for phylogenetic network reconstruction (Poormohammadi et al., 2020).

Computing information content of 3ts of a tree is problematic if they are not independent (i.e. absolute independence *sensu* Wilkinson et al., 2004). Nelson and Ladiges (1992) showed that, within a tree, the independence among a 3ts set could be achieved by using a fractional weighting to remove redundancy. Wilkinson et al. (2004) criticised this procedure by showing that specific relationships among 3ts generating dependency were not taken into account in the computation of fractional weighting. However, these relationships were neither clearly defined nor exhaustively stated in their work. A thorough work on relationships among 3ts in order to propose new operational solutions to compensate for the dependency becomes essential today. The goal of this paper is to develop our understanding about the relationship between dependency and 3ts inference rules that allows to deduce additional 3ts from a set of 3ts within a tree and to propose a weighting method to improve the accuracy of evolutionary analyses using 3ts.

In the first part of this paper, we summarise a historical review of the procedures proposed in the literature to compensate for the dependency in 3ts sets deduced from a tree. We highlight main challenges and discuss the solutions that have been proposed in the past. In the second section, we demonstrate the link between tree-shape and dependency within phylogenetic trees. We recognise different types of relationships between internal nodes that lead to dependency. In the third part, we exhaustively describe the different inference rules on 3ts and show their relationship to dependency. Our work on tree-shape and on 3ts is then used in a fourth part to propose a development of Nelson and Ladiges’ (1991b) procedure (herein called *corrected fractional weighting*) that removes all types of dependency between 3ts. This procedure is needed to fulfil the requirement of independence on which are based most of the analyses using phylogenetic information. Finally, we used simulated datasets to compare different supertree methods that use different representation of phylogenetic information and show that methods using fractional weighting performs significantly better.

## Dependency and Weighting 3ts: an historical review

The use of 3ts as a unit of measurement for the information content of a phylogenetic tree requires that they be independent of each other. Performing an analysis based on non-independent data leads to biases that can have important consequences on the quality of the result. Within a tree, the independence between two relationships implies that the truth of one does not involve the truth or falsity of the other. The 3ts *a*(*bc*) and *a*(*bd*) are independent because the truth of one tells nothing about the truth of the other. On the other hand, the 3ts *a*(*bc*), *a*(*bd*), and *a*(*cd*) are not independent relative to each other, because the truth of two of these 3ts necessarily implies the truth of the third (Nelson and Ladiges, 1992):

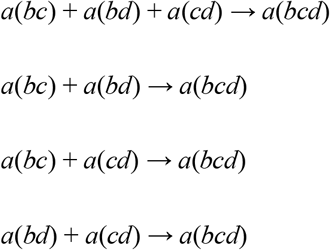

Relationships among 3ts may lead to dependency and therefore to redundancy. In the given example, the information content of these three 3ts cannot be 3; since the information given by the three 3ts is identical to that given by two of them, there is a repetition of information that leads to the overvaluation of 3ts.

Fractional weighting (FW; Nelson and Ladiges, 1992) was the first attempt to correct the overweighting of 3ts. Since any two 3ts are needed among the three, the amount of information carried by each 3ts is not one but 2/3. The corrected value of the set is 2. A component containing *t* taxa of which *n* are included in the informative node contains *n*(*t*-*n*)(*n*-1)/2 3ts, but only requires (*n*-1)(*t*-*n*) 3ts to be reconstructed. This quantity (*n*-1)(*t*-*n*) corresponds to the phylogenetic value of a component sensu Nelson and Ladiges (1992). In this case, the value of a 3ts is 2/*n*, i.e. the number of independent 3ts divided by the total number of 3ts.

The presence of several nested internal nodes in the tree implies another type of dependency. The first attempt of fractional weighting by Nelson and Ladiges (1992) was applied component by component, herein *fractional weighting per component* (FW_comp._) did not take into account the dependence. Indeed, in FW_comp._, the weight of a 3ts present in several components equals the sum of its weights for each component in which it appears. The tree *a*(*b*(*cd*)) can be decomposed into two components and then into 3ts:

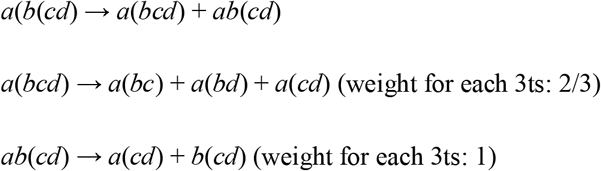

The information content of the tree is 4. 3ts weights are subject to redundancy: 3ts *a*(*cd*) is overweighted (5/3) because it is present in both components *a*(*bcd*) and *ab*(*cd*) while it is present only once in (*a*(*b*(*cd*))). For this reason, Nelson and Ladiges (1992: 491) describe FW_comp._ and *Uniform Weighting* (sum of 3ts per component, see table I) as “*both misleading*”. A 3ts cannot be present more than once in a tree, and therefore must not have a weight greater than 1. The table I shows another problematic example with the tree (*a*(*bc*(*de*))). To overcome this difficulty, Nelson and Ladiges (1992) propose a different way to calculate the absolute value of a 3ts called herein *total fractional weighting* (FW_total_). It is a reappraisal of the idea that the weight of a 3ts is the number of independent 3ts divided by the total number of 3ts, but at the scale of the whole tree, by summing the weights of 3ts for each component: “*For each of their cladograms, we list the number of independent three-taxon statements and the absolute value of each of all possible statements, stated as the ratio of independent statements to all possible statements*” (Nelson and Ladiges, 1992: 491). The FW_total_ becomes for a 3ts the sum of all independent 3ts for each component / the sum of all 3ts for each component. For C components deducible from a phylogenetic tree:

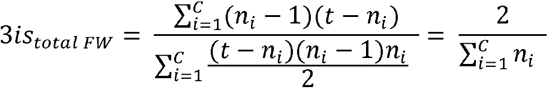

**Table I.**
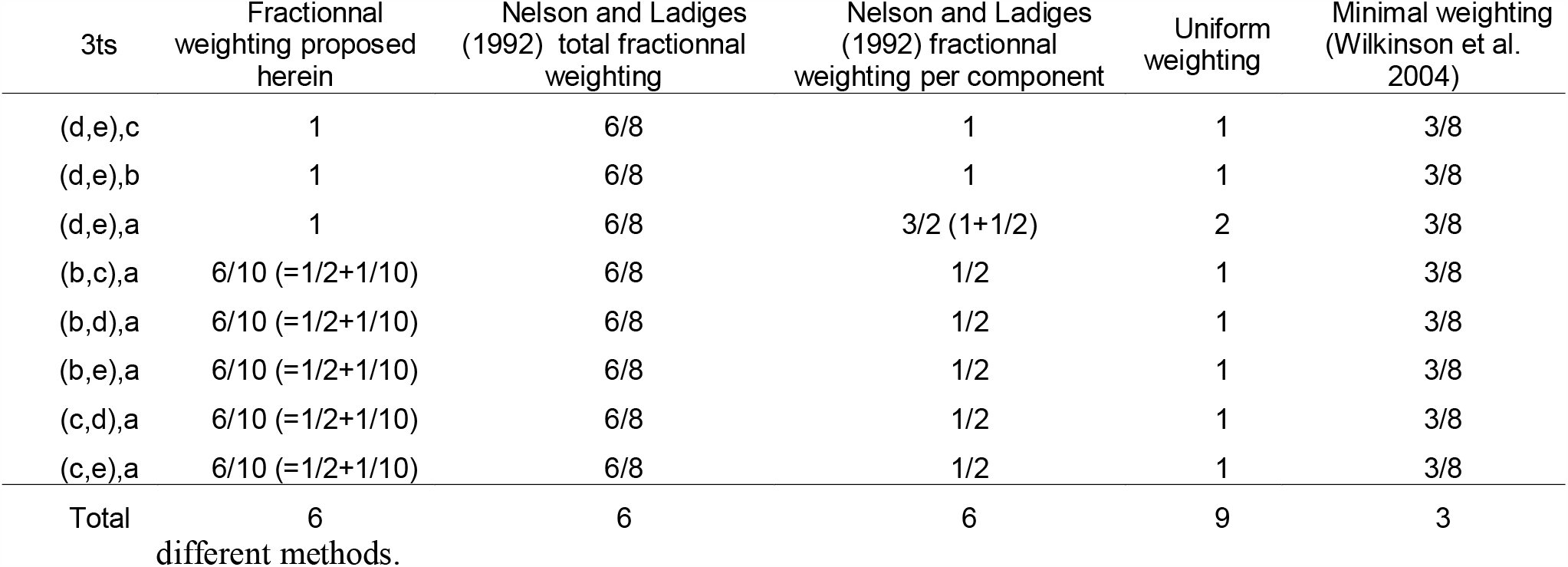
Decomposition of the tree (*a*(*bc*(*de*)) into 3ts and calculation of their weights using

| 3ts | Fractionnal<br>weighting proposed<br>herein | Nelson and Ladiges<br>(1992) total fractionnal<br>weighting | Nelson and Ladiges<br>(1992) fractionnal<br>weighting per component | Uniform<br>weighting | Minimal weighting<br>(Wilkinson et al.<br>2004) |
| --- | --- | --- | --- | --- | --- |
| (d,e),c | 1 | 6/8 | 1 | 1 | 3/8 |
| (d,e),b | 1 | 6/8 | 1 | 1 | 3/8 |
| (d,e),a | 1 | 6/8 | 3/2 (1+1/2) | 2 | 3/8 |
| (b,c),a | 6/10 (=1/2+1/10) | 6/8 | 1/2 | 1 | 3/8 |
| (b,d),a | 6/10 (=1/2+1/10) | 6/8 | 1/2 | 1 | 3/8 |
| (b,e),a | 6/10 (=1/2+1/10) | 6/8 | 1/2 | 1 | 3/8 |
| (c,d),a | 6/10 (=1/2+1/10) | 6/8 | 1/2 | 1 | 3/8 |
| (c,e),a | 6/10 (=1/2+1/10) | 6/8 | 1/2 | 1 | 3/8 |
| Total | 6 | 6 | 6 | 9 | 3 |
| different methods. |  |  |  |  |  |

As a result, all the 3ts of a tree have the same weight. The four 3ts of the tree *a*(*b*(*cd*)) have a weight of 1. Nelson and Ladiges conclude by considering this calculation of the absolute value of a 3ts as the most relevant (Nelson and Ladiges, 1992: 494).

Wilkinson et al. (2004) gave a new attempt to describe the dependencies between 3ts using Dekker’s (1986) dyadic inference rules. According to Dekker, only two inferences are possible from pairs of quartets to deduce additional quartets. By transposing these inference rules to 3ts, Wilkinson et al. (2004: 994) propose three rules for inferring 3ts exemplified as follows:

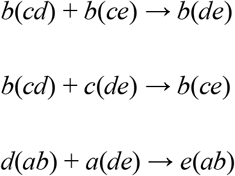

The original Dekker rules imply that any tree containing two quartets involved by the dyadic inference rules also contains an additional quartet. Wilkinson et al. (2004) point out that for 3ts, several rules can intervene from a single couple. For example :

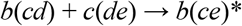

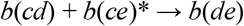

Which results in :

*b*(*cd*) + *c*(*de*) → *b*(*c*(*de*)) → *b*(*cd*) + *c*(*de*) + *b*(*ce*) + *b*(*de*) (fig. 4b in Wilkinson et al. 2004)

This implies that the presence of some couples of 3ts can imply the truth of two secondary 3ts. For Wilkinson et al. (2004), this type of dependence is not taken into account by Nelson and Ladiges’ fractional weighting. Since the tree (*b*(*c*(*de*))) needs only two 3ts to be rebuilt, its weight should be 2. Consequently, the FW_total_ can only take into account one type of dependence corresponding to the first rule presented above.

However, Wilkinson et al. do not give all the possible combinations that lead to additional 3ts. This is justified for them by the fact that any fully resolved tree with *t* taxa can be fully entailed by specific sets of *t*-2 3ts (Steel, 1992). Consequently, Wilkinson et al. believed that because any dichotomous tree includes only *t*-2 independent 3ts, each triplet of a tree can be weighted using the formula 6 / (*t* ^2^ -*t*). Wilkinson et al. named this procedure *Minimal Weighting* (MW), which aimed to take into account all possible inference rules without having to define them. The arguments put forward in favor of the MW are: (i) an identical weight for all 3ts because there is no argument to weight differentially the 3ts from a tree, (ii) a linear evolution of the total weight of a tree with the number of taxa (polynomial with FW_comp._ and FW_total_), and (iii) an evolution of the total weight of a tree that is not influenced by tree shape.

## Dependency and Tree Shape

Our aim being to find a way of deleting dependency issues in phylogenetic trees, we must define in a first place where the dependency can arise. Here we show for the first time how the tree shape can modify the dependency between nodes, and by corollary altering the information content of phylogenies. To fulfil this aim, we need to precisely define the various types of nodes of phylogenetic trees. There are three types of internal nodes: apical nodes, orthologous nodes, and paralogous nodes.

### Definition 1.

*Apical nodes* are nodes that only include leaves. The tree (*b*(*c*(*de*))) contains only one apical node, (*de*).

### Definition 2.

*Orthologous nodes* (sensu Zaragüeta et al., 2004, also called asymmetric or unbalanced nodes) include only one internal node as direct descendant. In the tree (*b*(*c*(*de*))), the node (*cde*) is orthologous because it leads only to the apical node (*de*), and the root node (*bcde*) because it immediately leads only to (*cde*).

### Definition 3.

*Paralogous nodes* (sensu Zaragüeta et al., 2004, also called symmetric or balanced nodes) are nodes that include two or more internal nodes as direct descendants. The tree ((*ab*)(*cd*)) contains one paralogous node, the root (*abcd*), which contains two internal nodes (*ab*) and (*cd*).

The relationships between internal nodes in the tree can be linked to the notion of dependence. Orthology implies dependency between nodes (a node is differentiated in another node), while paralogy implies independence between nodes, where a node is divided into several independent and separate lineages. We equate here dependency to inclusion; thus, dependency is a kind of information that may be taken into account, in contrast to independency (i.e. disjunction). Therefore, a pectinate tree contains more information that a symmetric tree containing disjointed independent nodes. As we believe as Williams and Ebach (2008: 213) that in a phylogenetic analysis, a multi-state character is more informative than the pair of corresponding binary characters (because the relationships between states are explicitly showed only in the multi-state character), we can say more broadly that a tree carries qualitatively a different information than the sum of all its components taken independently. The dependence or independence between nodes should have an impact on the information content, and thus on 3ts weights. Consequences for 3ts are explained in the next section.

## Three-taxon statement relationships and inference rules

Bryant and Steel (1995) have shown first that there are an infinite number of irreducible inference rules for 3ts (rules which are not deductible from each other). Subsequently, Rineau and Prin (2021) have proposed a new formalism of 3ts (Colonius and Shulze, 1981; McMorris and Powers, 2003) that resolves previous issues raised by Bryant and Steel: there is a finite list of 3ts inference rules that is enough to infer any other rule by repeated application. This finite list of inference rules is an essential tool to characterize how redundancy emerge within a set of 3ts coming from a tree, and thus to compensate for them. Moreover, because our aim here is to account for dependency rules in a 3is set coming from a *single tree*, the dyadic inference rules (i.e. when the combination of two 3is allow to deduce new 3is) stated below are necessary and sufficient to account for all cases of dependency, because specific cases shown in Rineau and Prin (2021:15-18) only arise from 3is coming from *several trees*.

We restate here the definitions of 3ts relationships responsible for dependency in order to show the link between 3ts inference rules, phylogenetic tree shape, and dependency. The specifics of relationships among 3ts depends on their common leaves and positions (Wilkinson et al., 2004; Rineau and Prin, 2021). Thus, two 3ts can be identical or not, compatible or not, combinable or not. The combinability relationship can itself be declined into four possible relationships from which are three inference rules allowing to generate additional 3ts (Rineau and Prin, 2021), and thus, redundancy.

### Definition 1.

Two distinct 3ts are *compatible* if they can be true at the same time, i.e. if there exists *at least* one tree containing both.

Consequently, two 3ts are incompatible if they cannot be true at the same time, i.e. the truth of one necessarily implies the falsity of the other. Two 3ts are incompatible if they have exactly the same taxa but state different relationships (e.g. *a*(*bc*) and *b*(*ac*)).

### Definition 2.

Two compatible 3ts are *combinable* if they can be unambiguously gathered in a single tree. In other words, two 3ts are combinable if and only if they are present in the strict consensus of all possible trees containing them. Two 3ts with their respective taxa sets D = {*u, v, w*} and E = {*x, y, z*} are combinable if there exists an informative strict consensus which taxa set *F* satisfies |F| = |D ⋃ E| = 4. We note that 3ts are combinable if and only if they have two taxa in common.

If two 3ts are not combinable, the above-mentioned strict consensus is a bush (a tree without any informative node). Combinability logically implies compatibility, but the reverse is not true: compatible couples of 3ts sharing less than two taxa are non-combinable. The couple *a*(*bc*) and *a*(*de*) is non-combinable because *a* is the only common taxon, and entails several possibilities, as ((*de*)(*a*(*bc*))) or (*a*(*b*(*d*(*ce*)))) and which strict consensus of all possible trees containing both 3ts is a bush (no unambiguous common statement between all possible trees).

Four relationships among combinable 3ts exist, with specific properties, depending on the location of their common taxa (Figure 1).

**Figure 1.**
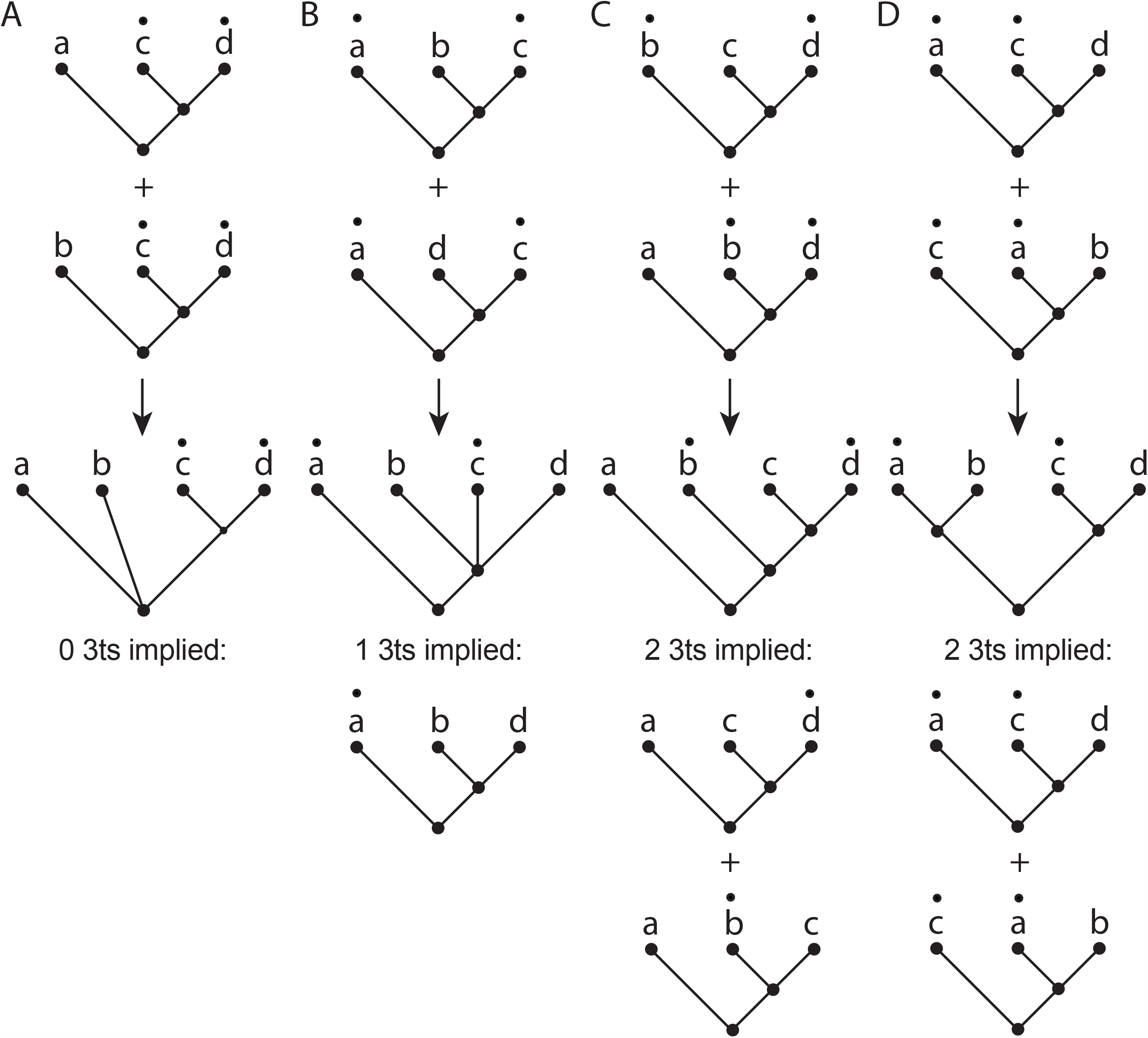
Four possible types of relationships between pairs of 3ts can be combined. Two 3ts sharing two taxa in common can be combined into a 4-taxon tree from which it is possible to deduct 0, 1 or 2 3ts. The black dots correspond to the leaves common to both 3ts. A. In-in nts relationship. B. In-out nts relationship. C. Asymmetric relationship. D. Symmetric relationship.

### Definition 3.

Two combinable 3ts are in *in-in nts relationship* if both common taxa are located inside the apical node of both 3ts, and the combination of the 3ts results in a single n-taxon statement (i.e. a tree with a single informative node; nts thereafter; Wilkinson, 1994).

The example of *a*(*cd*) + *b*(*cd*) allows us to build the tree *ab*(*cd*). The tree can be decomposed into the two 3ts that allowed its construction:

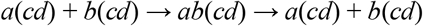

Information conveyed by each 3ts allows an additional taxon to be added as a sister group to the apical node (*cd*). The fact that no additional 3ts is generated implies that no dependency is generated: that is the reason two-leaf apical nodes never cause fractional weight, as Nelson and Ladiges (1992) noticed.

### Definition 4.

Two combinable 3ts are in *in-out nts relationship* when one of the common taxa is connected directly to the root and the other is connected to the apical node in both 3ts.

The combination of two 3ts in an in-out nts relationship leads to build a nts with a single informative node of three leaves.

The couple *a*(*bc*) and *a*(*cd*) illustrates the in-out nts relationship, whose analysis highlights the implication of a third 3ts (which makes it the first inference rule):

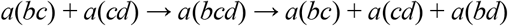

The in-out nts relationship between two 3ts generates a third 3ts with two taxa in common with the other two 3ts, at the same positions. The truth of these two 3ts implies the truth of a third. Among the three 3ts *a*(*bc*), *a*(*cd*), and *a*(*bd*), it can be seen that whatever the two 3ts chosen, they will be in the in-out nts relationship and will involve the third one. It is precisely this kind of relationship which is taken into account by Nelson and Ladiges’ FW to correct the redundancy. This type of dependency is the only one present within the components. If a new 3ts is added and forms additional in-out nts relationships, then all 3ts can be combined, as in-out relationship is transitive:

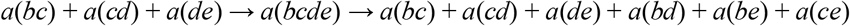

Three 3ts are linked by two in-out relationships. Each in-out relationship generates an additional 3ts:

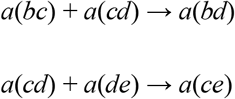

The generation of the secondary 3ts leads to the appearance of two new in-out dependency relationships, leading to the same 3ts:

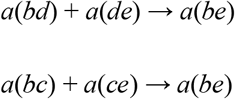

We call the initial 3ts as primary 3ts, *a*(*bc*), *a*(*cd*), and *a*(*de*) in the example. The 3ts generated from the combination of primary 3ts are called secondary 3ts, here *a*(*bd*) and *a*(*ce*). The 3ts generated by the combination of secondary 3ts are called tertiary 3ts, here *a*(*be*) and so on (to *n*-ary 3ts for the *n*th level of combination). For the combination of the set of three primary 3ts, a total of six *n*-ary 3ts are entailed. This is why coding a set of six 3ts *a*(*bc*), *a*(*cd*), *a*(*de*), *a*(*bd*), *a*(*be*), *a*(*ce*), requires a weight of ½ for each 3ts since only three 3ts among the six are needed to build the node (given that all terminals are represented among the set). From the three primary 3ts we can deduce all the secondary and tertiary 3ts. The corollary is that 3ts may not satisfy the pairwise compatibility theorem (Estabrook et al., 1976; Wilkinson, 1994b). Indeed, the fact that all primary 3ts are pairwise compatible does not imply that all n-ary 3ts are compatible.

### Definition 5.

Two combinable 3ts are in *asymmetric relationship* when both have a common taxon connected to the apical node, and when the other common taxon is connected to the apical node in one of the 3ts and to the root in the other.

Unlike with the in-in and in-out nts relationships, the trees constructed from two 3is linked by an asymmetric relationship have two informative nodes and an asymmetric shape (see fig. 4b in Wilkinson et al., 2004). If the tree (*a*(*b*(*cd*))) is decomposed into its two components *ab*(*cd*) and *a*(*bcd*), the first generates only one in-in nts relationship and the other three in-out nts relationships (Appendix 1A). The combination of these two components implies the existence of two new couples of 3ts that were not present in the independent components (second inference rule):

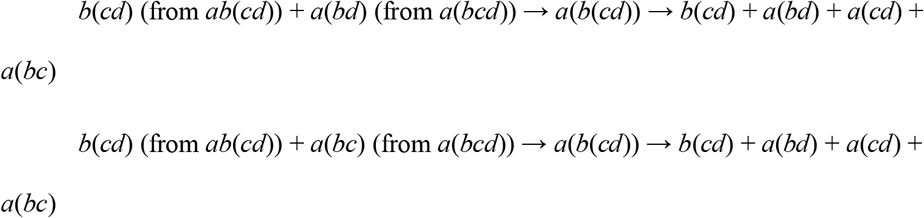

These two couples linked by an asymmetric relationship produce the complete tree *a*(*b*(*cd*)). If combined, 3is linked by asymmetric relationships, therefore, lead to inter-node dependency. This shows that the quantification of the information contained in a tree is not equal to the sum of the information of each component. Methods that decompose trees in their components without taking into account the relationships between these components, such as the supertree method MRP (Baum, 1992; Ragan, 1992) or the biogeographical method BPA (Wiley, 1986; Brooks, 1990), are thus flawed. The information conveyed by the inclusion (dependency) of one node in another must be taken into account.

### Definition 6.

Two combinable 3ts are in *symmetric relationship* when among the two taxa they have in common, one is connected to the root in the first 3ts and the apical node in the second 3ts, and the other is connected to the apical node in the first 3ts and to the root in the second. The symmetric relationship appears when at least two informative nodes are disjoint (i.e. if there is a paralogous node in the tree where the 3ts come from).

The following example illustrates a symmetric relationship between two 3ts that generates a symmetric tree (Appendix 1B):

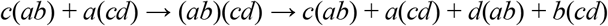

This combination allows two 3ts, *d*(*ab*) and *b*(*cd*), to be deduced. The symmetric relationship thus implies a third inference rule allowing to generate additional 3ts.

Some general corollaries may be listed from the set of above-described relationships:

– Any hierarchical tree with at least one informative node can be analysed into a set of 3ts without incompatible relationships.
– The definitions stated here are necessary and sufficient to cover all cases of dependency among a set of 3ts coming from a phylogenetic tree, as shown by Rineau and Prin (2021).
– Any tree containing an informative node and more than one leaf outside this informative node contains in-in relationships (Fig. 1A).
– Any tree containing an internal informative node and more than two leaves connected to that node contains in-out relationships (Fig. 1B).
– Compatible but not combinable 3ts (zero to one taxon in common) are present in a tree as long as there is an informative node comprising more than three leaves. As soon as there are at least two informative nodes in a tree, asymmetric (fig. 1C) and/or symmetric relationships (fig. 1D) appear. It is, therefore, possible to find all these types of relationships within the same tree (Appendix 1C).

## Weighting procedures

Formalised 3ts inference rules that lead to dependency are the only way to remove redundancy in 3ts decomposition. Firstly, we will show that MW fails to compensate for all instances of dependency. Secondly, we will propose an improvement of Nelson and Ladiges (1991b) procedure of fractional weighting based on the previously stated definitions that succeeds at removing every dependency arising from 3ts primary inference rules from any 3ts set.

### Minimal weighting

Wilkinson et al. (2004) designed the MW in order to remove the redundancy issues using a simple formula (*t*-2). However, there are two arguments for rejecting the MW.

The first concerns the identical weight given to each triplet, as in Nelson and Ladiges’ FW_total_ computation. This weight implies that among the four 3ts of (*b*(*c*(*de*))), only two are necessary to reconstruct the tree. The total weight of the tree (its information content) is 2, and each 3ts has an information content of ½. Wilkinson et al. (2004) rationale implies that each of the 3ts has the same information content. If this was true, any set of two 3ts among *b*(*cd*), *c*(*de*), *b*(*ce*), and *b*(*de*), would allow to reconstitute the tree, which is not correct. Here is the result of the different combinations of 3ts:

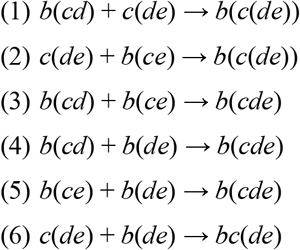

(1) and (2) are the only couples that allow the tree to be fully reconstituted. The other couples allow only one of the two components to be rebuilt: the couples (3) to (5) allow rebuilding *b*(*cde*), and the couple (6) allows rebuilding *bc*(*de*). In order to keep only the minimum number of 3ts, the statement “any 2 among 4” is incorrect. Therefore, MW is inconsistent with Wilkinson et al.’s (2004) rationale.

The second argument against MW is a strong limitation in its conception: it can only handle fully dichotomous trees. The total weight of a tree of *t* taxa is *t*-2 for a totally bifurcating tree and, to our knowledge, no analytical solution has so far been proposed to include multifurcations. Using the total weight formula of *t*-2 for polytomous trees implies that the information content of a given tree is identical to that of any of its subtrees as long as they have the same leaves. The trees ((*ab*)(*cd*)), (*ab*(*cd*)) and ((*ab*)*cd)* all have an information content of 2 according to this formula. This conclusion is clearly illogical.

### Corrected fractional weighting

To successfully eliminate dependency among a 3ts set deducted from a tree, the different types of 3ts inference rules (definitions 4 to 6) and the scale at which they operate (intra-node or inter-node) must be taken into account. MW fails to recognise in detail the different types of relationships. Both FW_total_ and FW_comp._ takes into account a single inference rule (in-out nts), which is insufficient. Our strategy to suppress the dependency is set up in sequential steps, each step corresponding to a type of 3ts inference rule (table II). We will explain here a corrected version of the FW (FW_cor._) that take into account all types of dependency defined above.

The first step corresponds to the procedure set out by Nelson and Ladiges in 1992. It consists in removing the dependency linked to in-out relationships within each component itself. The formula for the number of independent 3ts for a component is (*t*-*n*)(*n*-1), *t* being the total number of taxa and *n* being the number of taxa connected to the informative node (Nelson and Ladiges, 1992). The factor (*t*-*n*) corresponds to the number of leaves connected directly to the root. Increasing their number only generates in-in relationships. The factor (*n*-1) is linked to in-out relationships. For *a*(*bcde*), the factor (*n*-1) is 3. Only three 3ts are needed to rebuild the tree. The factor (*t*-*n*) is 1 and acts as a multiplier according to the number of leaves at the root. For example, by adding a leaf *f* to the root, (*n*-1) does not change. (*t*-*n*) on the other hand, equals 2:

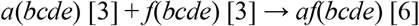

**Table II.**
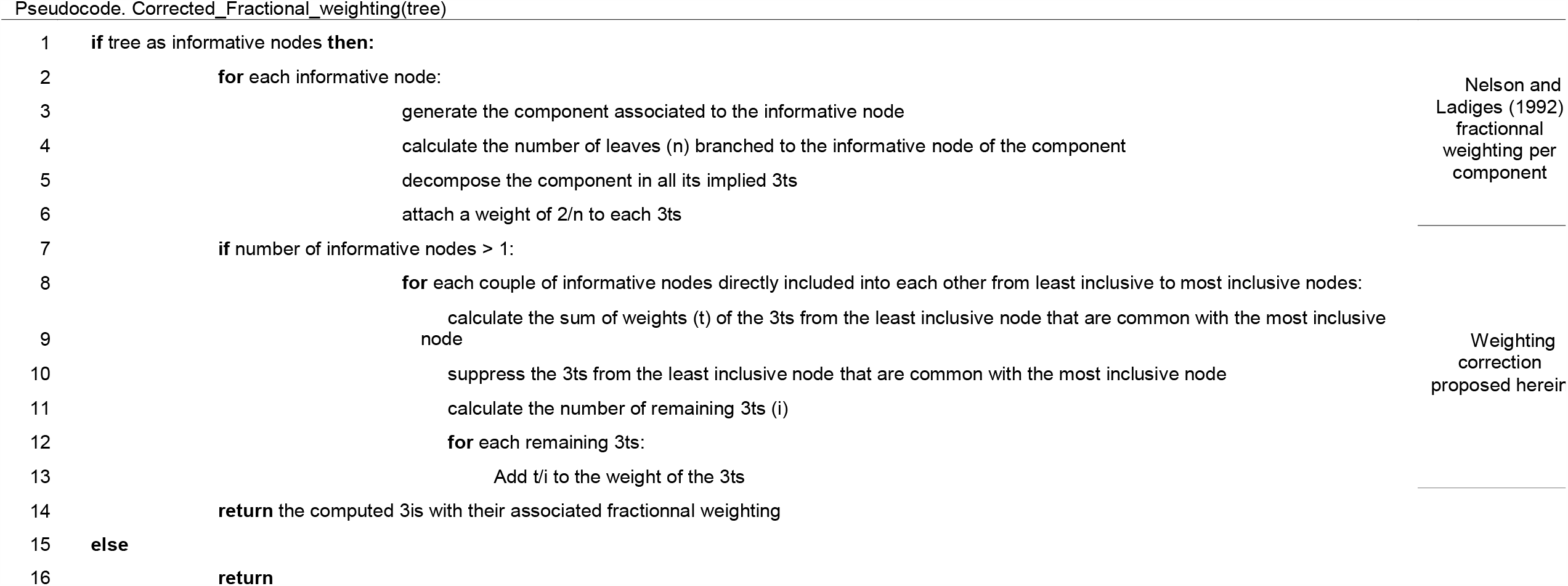
Pseudocode detailing the steps of removing the dependency of a set of 3ts from a tree by fractional weighting.

The numbers in square brackets are the weights of the corresponding trees. The minimum number of pairs of leaves pairs (e.g. *a,b* and *c,d*) necessary for each leaf to be in a pair and each pair to have a leaf in common with at least one another pair is (*n*-1). (*n*-1) is then multiplied by the number of leaves connected to the root (*t*-*n*) to give the formula (*t*-*n*)(*n*-1). (*t*-*n*) does not change between the formula of the number of independent 3ts and the formula of the total number of 3ts. On the other hand, the value (*n*-1) changes: to have the total number of 3ts, we need to search for all the combinations of two leaves among *n*, i.e.:

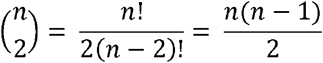

By replacing (*n*-1) by 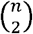 in (*t*-*n*)(*n*-1), we obtain the total number of 3ts:

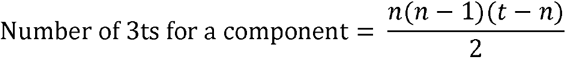

Nelson and Ladiges’ FW_comp._ is, therefore, FW_cor._’s first step in eliminating redundancy. This step of removing intra-node redundancy requires the analysis of each component independently. For each component, all its 3ts are generated, and the weight of 2/*n* (number of independent 3ts / total number of 3ts) is assigned to each 3ts. This weight eliminates overweight due to in-out relationships.

For example, component *a*(*bcd*) can be decomposed into three 3ts: *a*(*bc*), *a*(*cd*), *a*(*bd*). Each of the three pairs of 3ts forms an in-out nts relationship which logically implies the third 3ts. Each 3ts is therefore overweighed because it exists both as a primary and secondary 3ts. More generally, a 3ts present in a component is overweighed if it can be generated from other 3ts. This first step makes it possible to define the phylogenetic weight of each node.

FW_cor._ is obtained by removing the dependency linked to in-out nts relationships, as in FW_comp_, and then to asymmetric relationships among 3ts of different nodes (inter-node redundancy). Since 3ts have already been weighted in the previous step, the correction of the asymmetric dependency (i) concerns inclusions between informative nodes are present in the tree, and (ii) revises the weight of the 3ts assigned in the first step.

Nelson and Platnick (1991) showed that every 3ts generated from a fully pectinate tree has a weight of 1, applying FW or not because there is no redundancy between the 3ts of a fully pectinate tree. The correction of inter-node redundancy requires checking the 3ts of each pair of nodes involving direct inclusion. Two 3ts in an asymmetric relationship will generate secondary 3ts that already exist as primary ones. Using the example of the tree (*a*(*b*(*cd*))) analysed in Figure 2, Nelson and Ladiges (1992: 491) showed that the statement *a*(*cd*) occurs in both series. This 3ts is the only one having a weight greater than 1 5/3 (2/3 + 1): it is the overweighed 3ts. The component *a*(*bcd*) bears the redundant 3ts because it is the only component of the tree that contains this particular 3ts corrected by FW_comp_. The other component consists only of in-in relationships: its 3ts are therefore all essential to the reconstruction of the tree and have a weight of 1. The analysis of the two asymmetric relationships of the tree highlights the overweight of *a*(*cd*):

**Figure 2.**
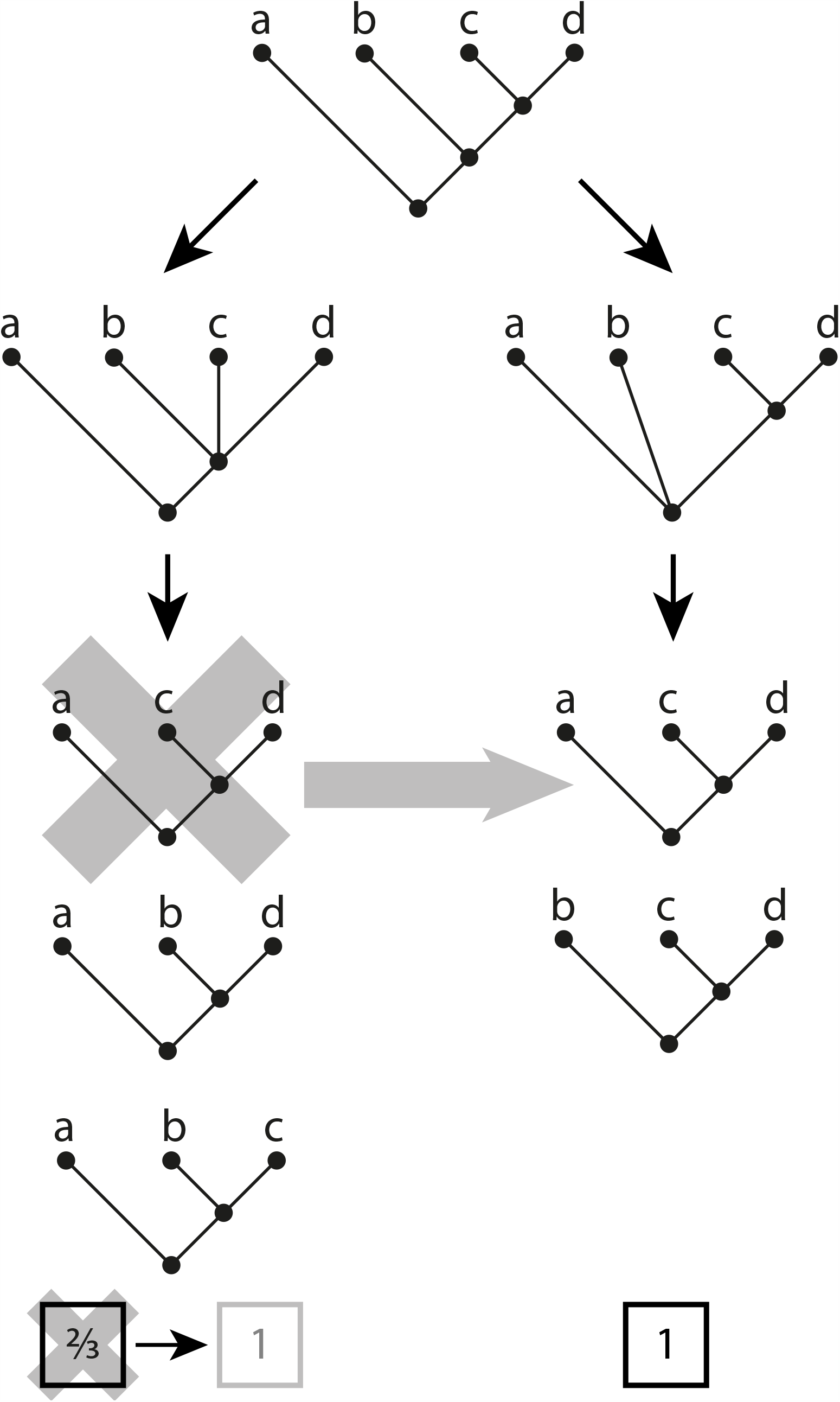
Fractional weighting procedure with correction of inter-node redundancy on (*a*(*b*(*cd*))).The dichotomous pectinate tree only produces 3ts of weight 1.

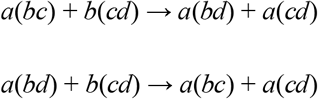

The secondary 3ts generated are *a*(*bd*), *a*(*bc*) and *a*(*cd*) (twice). All the 3ts resulting from asymmetric relationships of both components allow generating only one component: *a*(*bcd*). However, among all the 3ts, *a*(*cd*) appears twice while the others appear only once because the asymmetric relationship that overweighed this particular 3ts produces also a supplementary secondary 3ts, the only 3ts present in both components. The uncertainty of ‘two 3ts out of three’ from the component *a(bcd)* can then be resolved: the two necessary 3ts are *a*(*bd*) and *a*(*bc*). *a*(*cd*) is deleted from the component *a*(*bcd*), and its weight (2/3) is equally distributed among the others — if its weight was not distributed between the remaining 3ts, the result would be that knowledge about asymmetric relationship between nodes would lead to a loss of phylogenetic information. In (*a*(*b*(*cd*))), the weight of each of the four 3ts becomes 1. The ambiguity is resolved. The analysis of inter-node relationships allows us to correct the weights of 3ts from intra-node relationships because the inclusion of one node in another carries specific phylogenetic information.

To exemplify how polytomies are handled by this procedure, we apply these two steps to the tree (*a*(*bc*(*de*))) (Figure 3 and Table I). The six 3ts generated by the first component are weighted 1/2 (in-out relationships) and the three 3ts generated by the second component are weighed 1 (in-in relationships). The total information content of the tree is 6. It is noted that *a*(*de*) is present in both components, and can, therefore, be removed from the component *a*(*bcde*). After correction, the total weight of the tree is still 6, but the weights of the 3ts have been corrected. This tree shows a polytomy. Unlike the weights of the 3ts derived from the tree (*a*(*b*(*cd*))), which all end up balanced at 1, the weights of some of the 3ts derived from (*a*(*bc*(*de*))) are still fractional, even if the correction makes them tend towards 1. The dichotomous pectinate structure is the most informative: it contains the maximum number of components, and these components are all dependent on each other. Uncertainty increases with the number of polytomies, and with the number of branches involved in polytomies.

**Figure 3.**
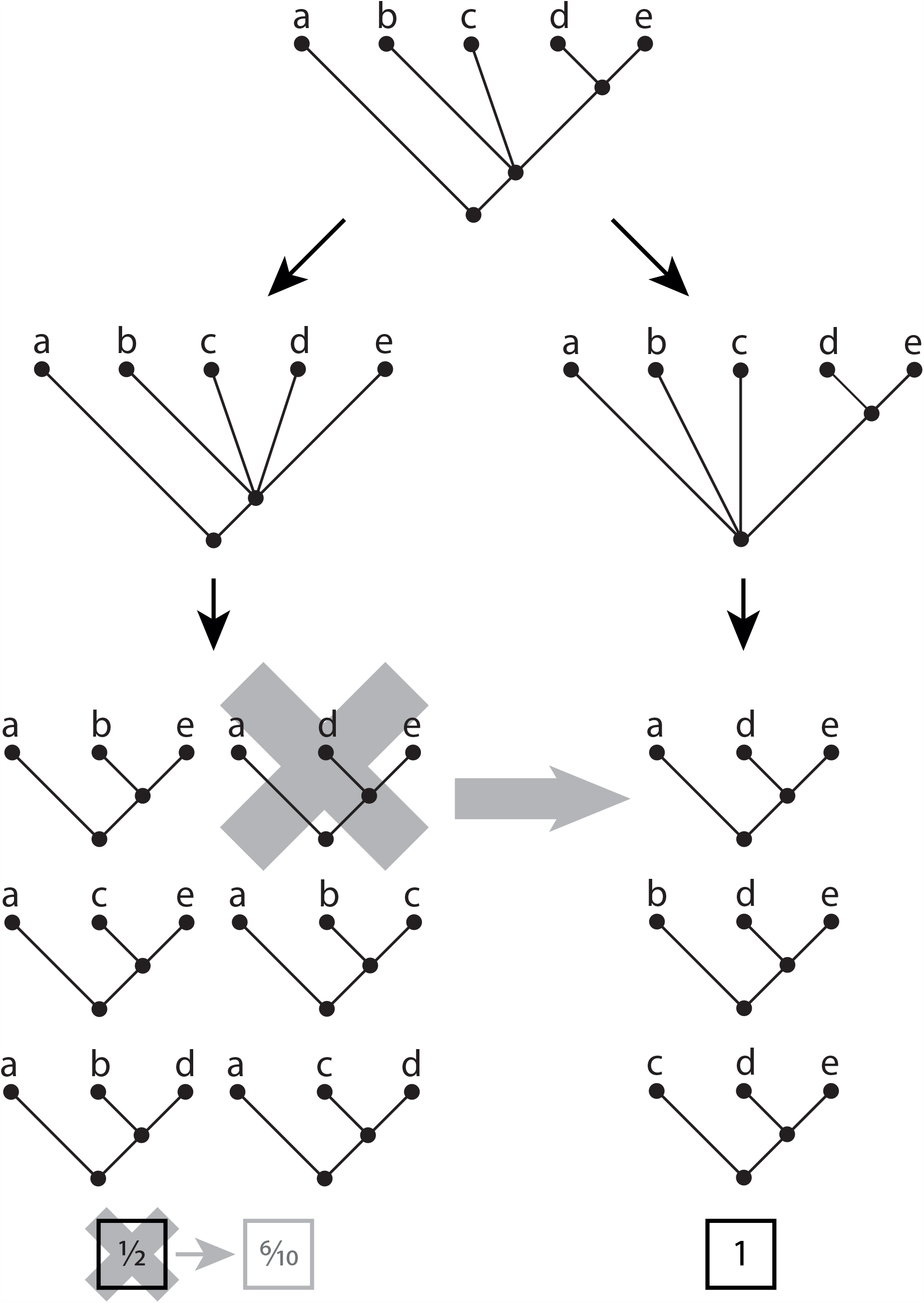
Fractional weighting procedure on (*a*(*bc*(*de*)). The tree is not completely resolved, which leads to uncertainty amongst 3ts and the persistence of fractional weightings for five of them.

The last type of dependency is related to symmetric relationships. Since symmetric relationships only concern independent, i.e. disjoint, groups, there is no dependency to manage at the level of 3ts for correcting 3ts weights. Indeed, symmetric relationships never provide redundancy of any particular 3ts.

An example with the analysis of the tree ((*ab*)(*cd*)) will illustrate this point (Appendix 1B):

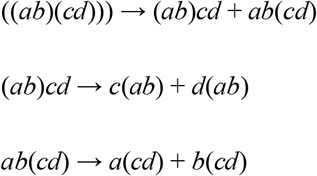

The weighting schemes proposed in the literature relay on the number of nodes. (i) The tree has a weight of 2 (corresponding to MW). Taken *independently*, each component of the tree also has a weight of 2; the choice of this weighting leads to accepting that the tree ((*ab*)(*cd*)) contains the same amount of phylogenetic information as (*ab*)*cd* or *ab*(*cd*). (ii) The tree has a weight of 4 and can be divided into two components without any loss of information (corresponding to the FW). However, since these two components are disjoint, they are independent. Here, the tree information is the sum of the information of its two independent components. Both components have a weight of 2. Symmetric relationships do not create differential weights between the 3ts because no 3ts is overrepresented by n-ary statements. Therefore, there is no justification for correcting in any way the weight of the 3ts intra-node.

Following Nelson and Ladiges (1992: 492): ‘*For cladogram 11, there is one internal node, ((AB)(CD)), with four statements, each with an absolute value of ¾*’. The authors provided no justification of their result (their result cannot be reached nor with the FW_comp._, nor with the FW_total_). However, as already pointed out by Nelson and Ladiges, if two totally dichotomous trees with the same number of leaves (more than four) are compared, the tree with more symmetric nodes will have less information content. The information content does need to reflect that. The information is maximal in pectinate trees because they are devoid of independent couples of nodes. For example, the weight of (*a*(*b*(*c*(*de*)))) is 10 whereas the weight of (*a*((*bc*)(*de*))) is 9.

To conclude, we would like to link the concept of phylogenetic information with the fractional weighting procedure. The literature on the concept of phylogenetic information was reviewed by Wilkinson et al. (2004). The authors concluded that a binary measurement of tree resolution was the most relevant measurement of information as defined by Shannon and Weaver. We disagree with their conclusion. We think that the idea of phylogenetic information cannot be isolated from the hypotheses proposed by a systematist to justify the characters that produce a particular tree. We thus link the idea of phylogenetic information to the notion of phylogenetic knowledge. The information-as-resolution can be measured on a tree. However, if the tree has been obtained by neighbor-joining, maximum likelihood or three-taxon analysis, it will not convey the same amount, or the same quality, of information. Moreover, this measurement fails to justify why a dichotomous tree should represent a maximally informative tree in terms of phylogenetic information. There is no theoretical or empirical justification that dichotomy is more or less phylogenetically informative than polytomies, except in terms of dichotomous resolution, which is circular.

The analytical procedure inherent to three-taxon analysis allows proposing a different measurement of phylogenetic information of a tree or a set of trees, as the sum of the fractional weighting of its/their three-item statements. This measurement allows to take into account the information conveyed by different characters that propose the same relationships among taxa. It seems obvious to as that having a particular node supported by several character-states may be linked to phylogenetic information, or to phylogenetic knowledge. We do not deny that the measurement proposed by Wilkinson et al. (2004) has some interest. However, the sum of the fractional weights of three-item statements seem to much better measure the information, or knowledge, that justifies the choice of a particular tree, i.e. the characters. It also allows to distinguish the information that identical trees may convey depending on the quantity and quality of characters and the theoretical background and method that was used to obtain it.

Our amended version FW_cor._ is the unique weighting scheme that takes into account dependency emerging both from intra-node (in-out relationships) and inter-node redundancy (asymmetric relationships), which leads us to believe that this is also a highly accurate way to quantify phylogenetic information in the cladistic paradigm. Our proposal of FW_cor._ has been implemented in the software LisBeth (Zaragüeta et al., 2012).

## Simulations

### Material and Methods

Following theoretical foundations of the relationship between dependency and weighting developed in the previous parts, our aim here is to simulate datasets and analyses to empirically test the efficiency of FW_cor._ versus current alternative methods. Each simulation consists in 1) generate a tree (named hereafter ‘the original tree’), 2) produce two subtrees, 3) generate noise in the subtrees, 4) use the subtrees with different weighting schemes to 5) reconstruct an optimal tree (and produce a strict consensus if several optimal trees exist) and compare this optimal tree (or the strict consensus of all optimal trees) with the original one to assess the relative efficiency of the different weighing schemes (Fig. 4). Each original tree is randomly generated using the python package ete3 (Huerta-Cepas et al., 2016) and contains 11 terminal taxa. Two subtrees are generated. Each is a copy of the original tree from which a defined number *m* of terminal taxa has been removed (*m* goes from 0 to 8; the choice of taxa is random). For each generation, 1000 simulations are run, leading to a total of 9000 runs. All the runs were repeated by introducing a varying amount of random noise by taxa permutations to simulate homoplasy and/or distortion. The different amount of distortion is included by pruning and regrafting a number *n* of taxa (*n* goes from 0 to 5). The choice of the pruned taxa and the regrafting locations are both random. The 9,000 runs were thus repeated 6 times leading to 54,000 runs. To combine both subtrees in the optimal tree, we used two supertree methods to reconstruct the optimal tree, one based on a parsimony approach (i.e. Matrix Representation with Parsimony) using PAUP 4.0a (Swofford, 2003), and the other based on a 3ta approach (Three-taxon analysis; Nelson and Platnick, 1991) with three different weighing schemes: FW_cor._, FW_comp._, and MW (in our simulations MW and FW_total._ give always the exact same results because the trees of each analysis always have the same number of terminal taxa), using Lisbeth (Zaragüeta et al., 2012). Note that when all trees of an analysis have the same number of taxa, the total fractional weight would lead to the same results as the minimal weight because the weights of all 3ts are the same for a given tree. The total number of runs is increased to 216,000. The ability of the methods to reconstruct the original tree is assessed by three complementary metrics. Triplet distance metrics (Tavares, 2018) were preferred to Robinson-Foulds, because the latter is strongly influenced by wildcard taxa. This is a common problem in supertree simulation analyses (Penny et al., 1982). Finally, the triplet distance metrics can be decomposed for our purpose in true resolutions (*TR*) and false resolutions (FR; Grand et al., 2013; Rineau et al., 2018).

**Figure 4.**
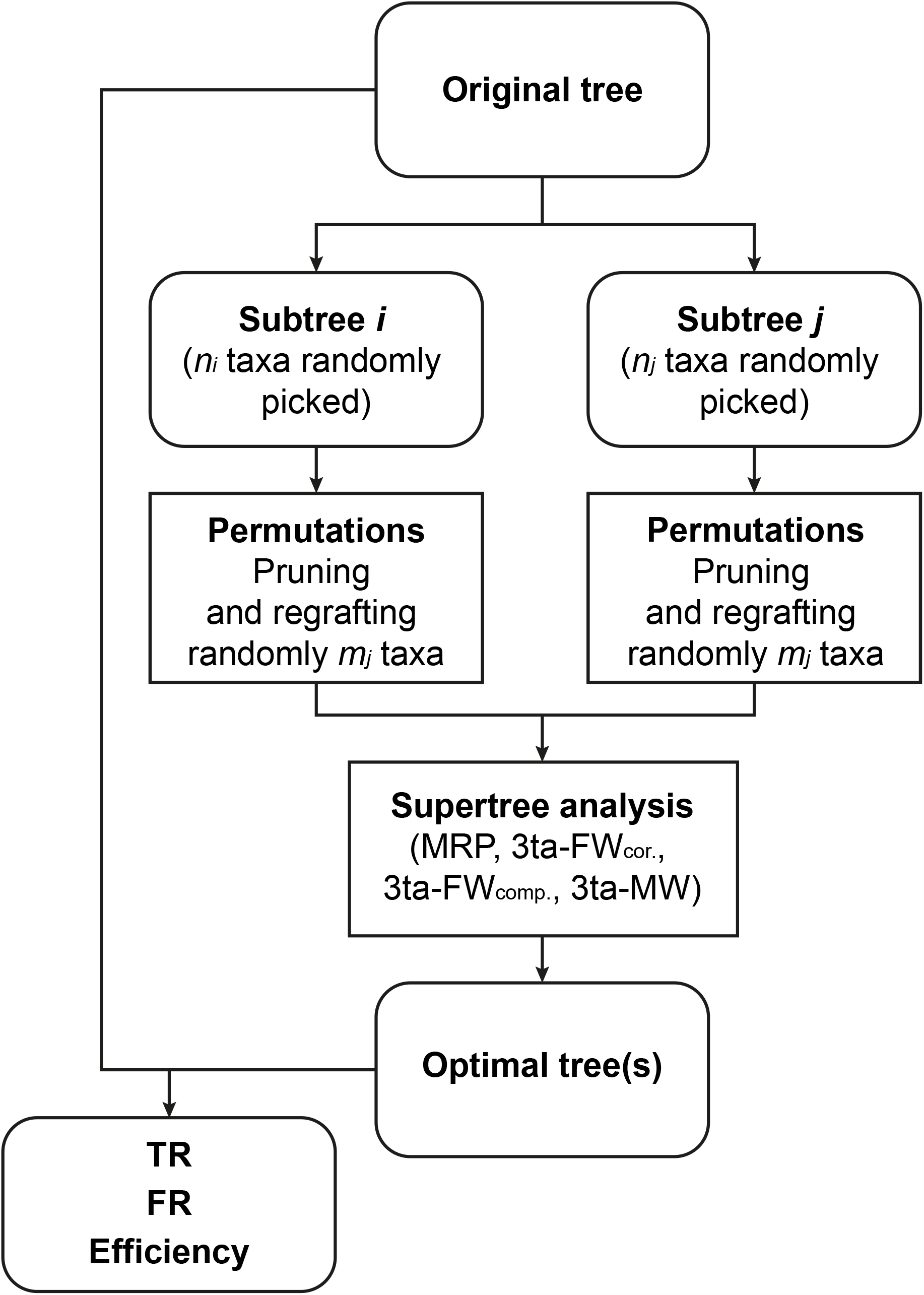
Flow-chart describing the steps for each run of the simulations: generation of an original tree, sampling of two subtrees with *n* taxa, pruning and regrafting of taxa, supertree analysis, comparison of the optimal tree(s) with the original tree in order to calculate *TR, FR*, and efficiency.

The first one, named *TR*, is the number of 3ts founded in both the original and optimal trees divided by the number of 3ts of the original tree. This can be viewed as the proportion of the 3ts from the original tree that is present in the optimal one. The second one, named *FR*, is the number of 3ts founded in the optimal tree but not in the original tree, divided by the number of 3ts of the optimal tree. This can be viewed as the proportion of the 3ts from the optimal tree that were not present in the original one. At last, the efficiency (*TR* – *FR*; range: [-100; 100]), allow to synthesise the results. −100 is the case where all phylogenetic relationships are false and the tree is completely resolved (i.e. dichotomous); 100 where all phylogenetic relationships are true and the tree is completely resolved. A null result means that there are as many true and false phylogenetic relationships, regardless of how well resolved the optimal tree is. In order to characterise the quantity of information brought by optimal trees, we counted internal nodes and 3ts from optimal tree for maximising precision, hereafter named *constrict resolution* and *constrict information content* respectively.

The results were then analysed using R software (R Development Core Team, 2008). We first explored the simulations by performing a PCA to provide a global overview of the covariations between the variables involved in the analyses. These variables consist in the starting parameters (i.e. number of permutations and number of taxa in subtrees), the number of optimal trees as well as their number of taxa, both the resolution and the information content of the strict consensus and finally, the 3 measures of the results quality (i.e. true resolutions, false resolutions and efficiency). Then, we describe in more details the relationships between the variables of interest. As those relationships show an important degree of complexity, we were not able to model them in a satisfactory manner. As a consequence, we used linear models to describe and test the relative efficiency of the methods and weighting schemes as well as other variables that can modify their relative efficiencies.

## Results and discussion

The two first axes of the PCA represent 73 % of the dataset variance. The first axis represents mostly the contribution of TR and four positively correlated variables that are: the constrict resolution, the constrict information content, the number *n* of taxa in subtrees, and the number of taxa in the optimal tree (related to the more or less important overlapping of the taxa sets of the subtrees; Fig. 5). For a given number of permutations, the amount of TR compared to the number of taxa in subtrees has an S-shaped distribution (see Fig. S1A), consequently TR tends to a horizontal asymptote (e.g. 100% of TR for 0 permutations when the number of taxa in subtrees tends to the one in the original tree). As the number of permutations increases, the asymptote decreases. The second axis of the PCA is mostly structured by FR (Fig. 5). As expected, the number of permutations (i.e. the quantity of noise) correlates mainly with this axis. In details, for a given number of permutations, the number of FR firstly increases with the number of taxa in subtrees and then decreases (Fig. S1B). Obviously, the efficiency (TR – FR) positively correlates to the first axis (thus to TR) and negatively to the second axis (thus to FR) (Figs. 5, S1C). Finally, the number of optimal trees does not correlate much with the other variables.

**Figure 5.**
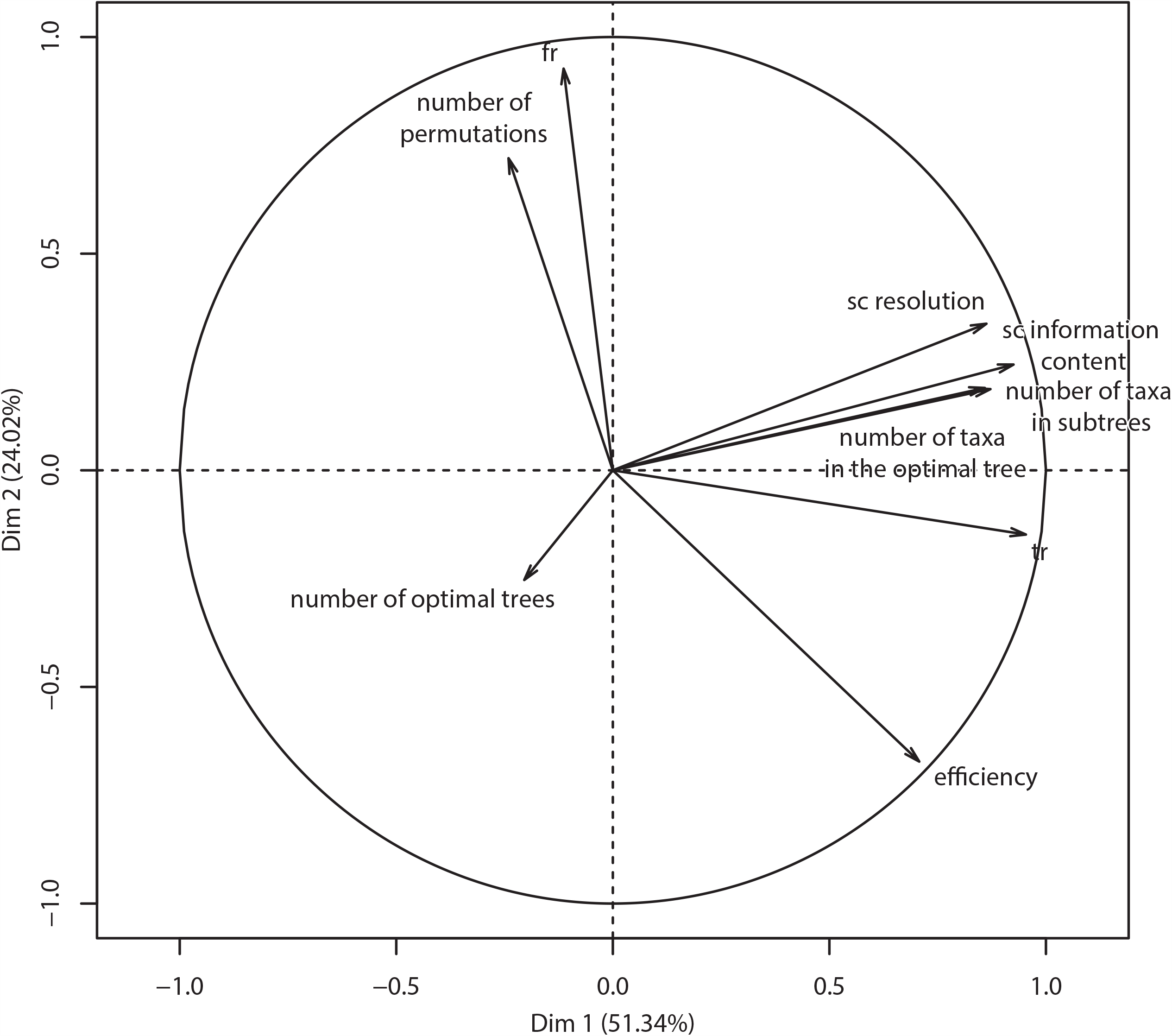
Variables factor map of the PCA for the two first principal components (*PCA* function from *FactoMineR* R package). The first axis represents 51.34% of the variance and correlates positively with *TR* (true resolutions), numbers of taxa in subtrees and optimal tree, strict consensus (sc) information content and resolution. The second axis represents 21.02% of the variance and correlates positively with FR (false resolutions) and the number of permutations. The variance of the efficiency is split between the first and second axis.

The relationships between all variables were thoroughly explored. The relative amount of TR and FR depends also greatly on the number of taxa in the optimal tree (Fig. S2). For a given number of taxa in the optimal tree, most of the runs fall on a line going from 0% TR and 100% FR to 0% FR and a percentage *p* of TR. This percentage *p* reaches 100% when the number of taxa in the optimal tree equals the one in the original tree. These lines represent the maximal resolution reachable with fully bifurcating optimal trees.

As concluded from the PCA, the variance of the dataset can be mostly summed up by TR, FR, the number of taxa in subtrees, and the number of permutations. The relationships between those variables are mostly monotonous. Thus, we computed two linear models with respectively TR and FR as dependent variables and as independent variables for both models: the number of taxa in subtrees, the number of permutations, and the different types of analyses (MRP, 3ta with FW_cor._, FW_comp., or_ MW/ FW_total_). These models thus have three variables and we included all the interactions terms. The homoscedasticity is not verified. For example, the variance of TR drastically increases in relation to the number of taxa in subtrees. As a consequence, we do not expect from these models to test for the effects but rather to investigate main tendencies and to observe specific interactions between the different types of analyses and the other variables (summaries of linear models are provided in Table S2). We use the function *interact_plot* from the R package *jtools* to visualise to three interactions as well as main effects from isolated variables at the same time (Fig. 6). Linearity checks are provided in the supplementary information (Fig. S3). The relationships are consistent with what inferred from the PCA. The increase of TR with the number of taxa in subtrees depends on the number of permutations. The higher the number of permutations is, the lower the increase of TR in relation to the number of taxa in subtrees. The main results with the substantial effect are that FW_cor._ and FW_comp._ are indistinct from each other and better compared to MRP and MW as the number of taxa in subtrees increases. The increase of FR with the number of permutations also depends on the number of taxa in subtrees and the type of analysis. For a few taxa in subtrees, the analyses are indistinct no matter the number of permutations. With both the increase of the number of taxa and permutations, differences between analyses increase with the order of FR amount: MW, MRP, and indistinctively FW_cor._ and FW_comp._ (Fig. 6). The efficiency decreases with the number of permutations. It also increases with the number of taxa in subtrees, especially at a low number of taxa. The numerous taxa in subtrees are, the higher the slope of this relationship is. In other terms, the efficiency does not really increase with the number of taxa in subtrees for too noisy datasets. This being said, FW_cor._ and FW_comp._ have better efficiencies compared to MRP and MW as the number of taxa in subtrees increases (Fig. 6).

**Figure 6.**
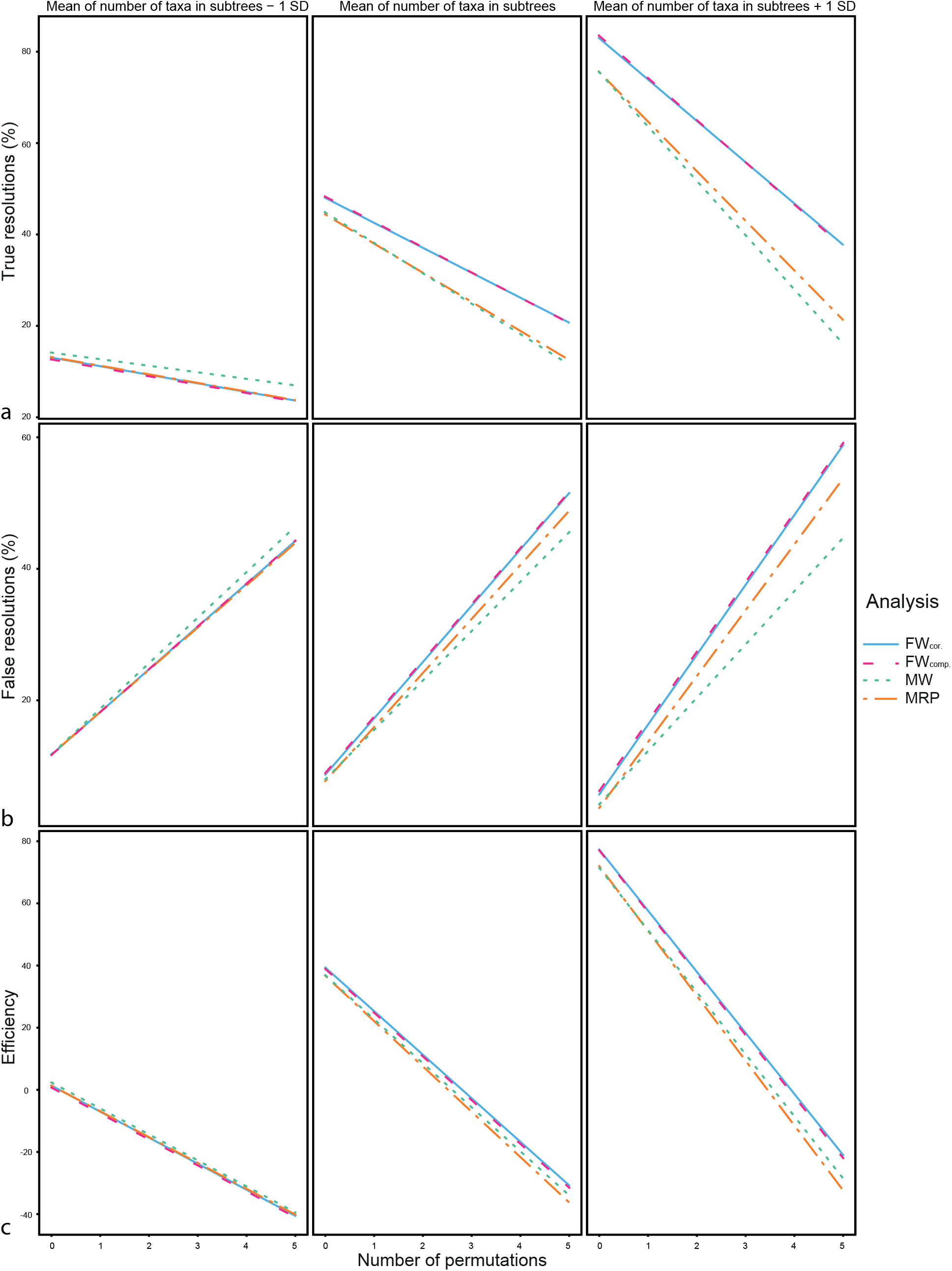
Interaction plots illustrating three linear models all including as independent variables the number of taxa in subtrees, the number of permutations and the types of analysis (function *interact_plot* from *jtools* R package. a. TR (number of true resolutions) as dependent variable. b. FR (number of false resolutions) as dependent variable. c. Efficiency as dependent variable. Types of lines represent the types of analysis, solid line: FW_cor._, dashed line: FW_comp._, pointed line: MW, dashed and pointed line: MRP.

In conclusion, FW_cor._ and FW_comp._ provide more resolved trees in comparison to MRP and MW. As the number of true resolutions (TR) is generally higher than the number of false resolutions (FR), efficiency is a bit better for FW_cor._ and FW_comp_. This leads us to recommend these weighting methods for the purpose of supertree reconstruction methods, cladistics methods, or metrics using 3ts. These are very preliminary results and they need to be confirmed in other conditions such as the type of noise, the tree shape. Future works will have to study more precisely the differential effect of FW_cor._ versus FW_comp._, which was indistinguishable in our simulations. The absence of significant differences between FW_cor._ and FW_comp._ may be related to the very low number of source trees. The higher the number of source trees is, the more incongruent relationships there is. It is the incongruence that will lead to the choice of this or that 3ts, and the choice should be biased with the FW_comp._ which will weight some 3ts to more than 1, which is theoretically flawed. The most obvious lead to explain these results could be the parameters of the analysis.

## Conclusion

Our paper focuses on the different types of relationships within a 3ts set coming from a phylogeny, some of them being directly responsible for the emergence of dependency. We also proposed a method for evolutionary biologists whose aim is to completely eliminate dependency from which all methods based on three-taxon statements currently suffer.

We propose for the first time a method of 3ts weighting based on intra-node and inter-node redundancy. Ideally, any phylogenetic or supertree analysis should only include datasets whose elements (hierarchical trees) are independent of each other, to eliminate any possibility of bias. We have shown that the simple decomposition of a tree into its components leads to a loss of information and that our method of corrected fractional weighting is the only procedure able to fully delete all the redundancies. We have also shown using computer simulations that the absence of weighting can significantly reduce the effectiveness of any method based on the 3ts. We also proposed the first empirical comparison between supertree component methods (MRP) and 3ts (three-taxon analysis), confirming the efficiency of methods which decompose trees into 3ts. More precisely, the most efficient methods are those which weight 3ts through a decomposition step into components, as Fractional Weighting. Finally, understanding the dependency relationships between the 3ts will permit to improve the accuracy and efficiency of supertree methods, consensus methods and phylogenetic analysis methods based on 3ts. “Triplet distance” metrics should expect the same handling. To conclude, we hope to open new debates and to fuel methodological discussions on how phylogenetic information should be weighted following 3ts inference rules in order to improve the accuracy of analyses.

## Acknowledgements

Valentin Rineau and René Zaragüeta conceived the ideas, Valentin Rineau acquired and analysed the datasets, and led the writing of the manuscript, Jérémie Bardin conducted the statistical analyses. All authors contributed critically to the drafts and gave final approval for publication. We deeply thank David Williams and Evgeny Mavrodiev for their fruitful comments on previous drafts, and to Stephane Prin and Paul Zaharias for discussions on 3ts combination procedures. We also thank Stephane Prin, Paul Zaharias, and Rachel Vautrin for comments on the draft and corrections. This work has been founded by the UMR 7207, Centre de Recherche en Paléontologie – Paris, France.

## Supporting information

Figure S1.

Boxplots of true resolutions (TR), false resolutions (FR) and efficiency in regard to the number of taxa in subtrees, split by the type of analysis (MRP: matrix representation with parsimony, FW_cor._: 3ta with corrected fractional weighting, MW: 3ta with minimal weighting, FW_comp._: 3ta with fractional weighting per component) and the number of permutations.

Figure S2.

Movie of a three dimensions block diagram with the percentages of true and false resolutions (TR and FR), the number of taxa in the optimal tree (also highlighted by colors). Darkening of each color depends on the resolutions of the optimal tree, dark corresponds to poorly resolved trees and bright corresponds to well-resolved tree.

Figure S3.

Loess smoothed lines indicating a deviation from linearity effect from the three linear models.

Table S1

A. Analysis of the different types of relationships between 3ts from a pectinate tree (*a*(*b*(*cd*))) and its components *ab*(*cd*) and *a*(*bcd*) taken independently. B. Analysis of the different types of relationships between 3ts from a symmetric tree ((*ab*)(*cd*)) and its components *ab*(*cd*) and (*ab*)*cd* taken independently. C. Analysis of the different types of relationships between 3ts from a mixed tree (((*ab*)(*cd*))*e*) and its components (*ab*)*cde, ab*(*cd*)*e*, and (*abcd*)*e* taken independently. The different types of relationships are in-in, in-out, asymmetric (a), symmetric (s). An empty cell means 3ts are compatible but not combinable.

Table S2.

Summaries of the three linear models containing formulas of the models (bold), the estimates of each coefficient as well as standard error, t-values, and p-values.

## Literature cited

Adams EN. 1986. N-trees as nestings: Complexity, similarity, and consensus. Journal of Classification 3:299–317.

Aho AV, Sagiv Y, Szymanski TG, Ullman JD. 1981. Inferring a tree from lowest common ancestors with an application to the optimisation of relational expressions. SIAM Journal on Computing 10:405–421.

Baum B. 1992. Combining Trees as a Way of Combining Data Sets for Phylogenetic Inference, and the Desirability of Combining Gene Trees. Taxon 41:3–10.

Brooks DR. 1990. Parsimony Analysis in Historical Biogeography and Coevolution: Methodological and Theoretical Update. Systematic Zoology 39:14–30.

Bryant D. 2003. A classification of consensus methods for phylogenetics. DIMACS series in discrete mathematics and theoretical computer science 61:163–184.

Bryant D, Steel M. 1995. Extension Operations on Sets of Leaf-Labelled Trees. Advances in applied mathematics 16:425–453.

Cao N, Bourdon E, El Azawi M, Zaragüeta R. 2009. Three-item analysis and parsimony, intersection tree and strict consensus: a biogeographical example. Bulletins de la société géologique de France 180:13–15.

Colonius H, Schulze HH. 1981. Tree structures for proximity data. British Journal of Mathematical and Statistical Psychology 34:167–180.

Dannenberg K, Jansson J, Lingas A, Lundell EM. 2019. The approximability of maximum rooted triplets consistency with fan triplets and forbidden triplets. Discrete Applied Mathematics 257:101–114.

Dekker MCH. 1986. Reconstruction methods for derivation trees. Unpublished Master Thesis, Vrije Universiteit.

Estabrook GF, Johnson JCS, McMorris FR. 1976. A mathematical foundation for the analysis of cladistic character compatibility. Mathematical Biosciences 29:181–187.

Farris JS. 1970. Methods for Computing Wagner Trees. Systematic Zoology 19:83–92.

Fitch WM. 1970. Distinguishing homologous from analogous proteins. Systematic Zoology 19:99–113.

Grand A, Corvez A, Duque Velez LM, Laurin M. 2013. Phylogenetic inference using discrete characters: performance of ordered and unordered parsimony and of three-item statements. Biological Journal of the Linnean Society 110:914–930.

Huerta-Cepas J, Serra F, Bork P. 2016. ETE 3: Reconstruction, Analysis, and Visualization of Phylogenomic Data. Molecular Biology and Evolution 33:1635–1638.

Islam M, Sarker K, Das T, Reaz R, Bayzid MS. 2020. STELAR: A statistically consistent coalescent-based species tree estimation method by maximising triplet consistency. BMC genomics 21: 1–13.

Kitching IJ, Forey PL, Humphries CJ, Williams DM. 1998. Cladistics: The Theory and Practice of Parsimony Analysis. Second edition. Oxford: Oxford University Press.

Kuhner MK, Yamato J. 2015. Practical Performance of Tree Comparison Metrics. Systematic Biology 64:205–214.

Liu L, Yu L, Edwards SV. 2010. A maximum pseudo-likelihood approach for estimating species trees under the coalescent model. BMC Evolutionary Biology 302:1–18.

McMorris FR, Powers RC. 2003. The Arrovian Program from weak orders to hierarchical and tree-like relations. In: Janowitz MF, Lapointe FJ, McMorris FR, Mirkin B, Robert FS, eds. Bioconsensus. Providence: American Mathematical Society. DIMACS series in discrete mathematics and theoretical computer sciences 61:37–45.

Mickevich MF, Platnick NI. 1989. On the information content of classifications. Cladistics 5:33–47.

Nelson G. 1979. Cladistic analysis and synthesis: principles and definitions, with a historical note on Adanson’s Familles des Plantes (1763–1764). Systematic Biology 28:1–21.

Nelson G, Ladiges PY. 1991a. Standard assumptions for biogeographic analysis. Australian Systematic Botany 4:41–58.

Nelson G, Ladiges PY. 1991b. Three-area statements: standard assumptions for biogeographic analysis. Systematic Biology 40:470–485.

Nelson G, Ladiges PY. 1992. Information content and fractional weight of three-item statements. Systematic biology 41:490–494.

Nelson G, Ladiges PY. 1994. Three-item consensus empirical test of fractional weighting. Systematics Association Special Volume 52:193–209.

Nelson G, Platnick NI. 1981. Systematics and Biogeography: Cladistics and Vicariance. New York: Columbia University Press.

Nelson G, Platnick NI. 1991. Three-Taxon Statements: A More Precise Use of Parsimony? Cladistics 7:351–366.

Penny D, Foulds LR, Hendy MD. 1982. Testing the theory of evolution by comparing phylogenetic trees constructed from five different protein sequences. Nature 297:197–200.

Poormohammadi H, Zarchi MS, Ghaneai H. 2020. NCHB: A Method for Constructing Rooted Phylogenetic Networks from Rooted Triplets based on Height Function and Binarization. Journal of Theoretical Biology 2–35.

Prin S. 2012. Structure mathématique des hypothèses cladistiques et conséquences pour la phylogénie et l’évolution. Avec une perspective sur l’analyse cladistique. Unpublished PhD Thesis, Muséum National d’Histoire Naturelle.

R Development Core Team. 2008. R: A language and environment for statistical computing. Vienna: R Foundation for Statistical Computing.

Ragan M. 1992. Phylogenetic Inference Based on Matrix Representation of Trees. Molecular Phylogenetics and Evolution 1:53–58.

Ranwez V, Criscuolo A, Douzery EJP. 2010. SUPERTRIPLETS: a triplet-based supertree approach to phylogenomics. Bioinformatics 26:i115–i123.

Rineau V, Zaragüeta R, Laurin M. 2018. Impact of errors on cladistic inference: simulation-based comparison between parsimony and three-taxon analysis. Contributions to Zoology 87:25–40.

Rineau V, Prin S. 2021. Cladistic hypotheses as degree of equivalence relational structure, and what it implies on three-item statements. bioRxiv: 2021.01.14.426769.

Sevillya G, Frenkel Z, Snir S. 2016. Triplet MaxCut: a new toolkit for rooted supertree. Methods in Ecology and Evolution 7:1359–1365.

Strimmer K, von Haeseler A. 1996. Quartet Puzzling: A Quartet Maximum-Likelihood Method for Reconstructing Tree Topologies. Molecular Biology and Evolution 13:964–969.

Swofford DL. 2003. PAUP*. Phylogenetic Analysis Using Parsimony (*and Other Methods). Version 4. Sunderland: Sinauer Associates.

Tavares BL. 2018. A synopsis of comparative metrics for classifications. 1804.03929.:37.

Vach W. 1994. Preserving consensus hierarchies. Journal of Classification 11:59–77.

Wiley EO. 1986. Methods in vicariance biogeography. In: Hovenkamp P, ed. Systematics and Evolution: a matter of Diversity. Utrecht: University of Utrecht Press, 283–306.

Wilkinson M. 1994a. Common Cladistic Information and its Consensus Representation: Reduced Adams and Reduced Cladistic Consensus Trees and Profiles. Systematic Biology 43:343–368.

Wilkinson M. 1994b. Three-taxon statements: when is a parsimony analysis also a clique analysis? Cladistics 10:221–223.

Wilkinson M, Cotton J, Thorley J. 2004. The Information Content of Trees and Their Matrix Representations. Systematic Biology 53:989–1001.

Williams DM. 2004. Supertrees, components and three-item data. In: O Bininda-Emonds, ed. Phylogenetic supertrees: combining information to reveal the tree of life. Dordrecht: Kluwer academic, 389–408.

Williams DM, Ebach MC. 2008. Foundations of systematics and biogeography. Dordrecht: Kluwer academic.

Williams DM, Humphries CJ. 2003. Component Coding, Three-item Coding, and Consensus Methods. Systematic Biology 52:255–259.

Zaragüeta R, Lelièvre H, Tassy P. 2004. Temporal paralogy, cladograms, and the quality of the fossil record. Geodiversitas 26:381–389.

Zaragüeta R, Ung V, Grand A, Vignes-Lebbe R., Cao N., Ducasse J. 2012. LisBeth: New cladistics for phylogenetics and biogeography. Comptes Rendus Palevol 11:563–566.

