## Supplementary material for "Information Content of Trees: Three-taxon Statements Inference Rules and Dependency": Table S1

Appendix 1

a. Analysis of the different types of relationships between 3ts from a pectinate tree (*a*(*b*(*cd*))) and its components (*ab*(*cd*)) and (*a*(*b*(*cd*)) taken independently. b. Analysis of the different types of relationships between 3ts from a symmetric tree ((*ab*)(*cd*)) and its components (*ab*(*cd*)) and ((*ab*)*cd*) taken independently. c. Analysis of the different types of relationships between 3ts from a mixed tree (((*ab*)(*cd*))*e*) and its components ((*ab*)*cde*), (*ab*(*cd*)*e*), and ((*abcd*)*e*) taken independently. The different types of relationships are in-in, in-out, asymmetric (a), symmetric (s). An empty cell means 3ts are compatible but not combinable.

a.

| Component 1: (a,b,(c,d)) | | |
| --- | --- | --- |
|  | b(c,d) | a(c,d) |
| b(c,d) | . | in-in |
| a(c,d) |  | . |

| Component 2: (a,(b,c,d)) | | | |
| --- | --- | --- | --- |
|  | a(b,d) | a(c,d) | a(b,c) |
| a(b,d) | . | in-out | in-out |
| a(c,d) |  | . | in-out |
| a(b,c) |  |  | . |

| Tree: (a,(b,(c,d))) (component 1 + 2) | | | | |
| --- | --- | --- | --- | --- |
|  | b(c,d) | a(b,d) | a(c,d) | a(b,c) |
| b(c,d) | . | a | in-in | a |
| a(b,d) |  | . | in-out | in-out |
| a(c,d) |  |  | . | in-out |
| a(b,c) |  |  |  | . |

b.

| Component 1: (a,b,(c,d)) | | |
| --- | --- | --- |
|  | a(c,d) | b(c,d) |
| a(c,d) | . | in-in |
| b(c,d) |  | . |

| Component 2: ((a,b),c,d) | | |
| --- | --- | --- |
|  | c(a,b) | d(a,b) |
| c(a,b) | . | in-in |
| d(a,b) |  | . |

| Tree: ((a,b),(c,d)) (component 1 + 2) | | | | |
| --- | --- | --- | --- | --- |
|  | c(a,b) | a(c,d) | d(a,b) | b(c,d) |
| c(a,b) | . | s | in-in | s |
| a(c,d) |  | . | s | in-in |
| d(a,b) |  |  | . | s |
| b(c,d) |  |  |  | . |

c.

| Component 1: ((a,b),c,d,e) | | | |
| --- | --- | --- | --- |
|  | c(a,b) | d(a,b) | e(a,b) |
| c(a,b) | . | in-in | in-in |
| d(a,b) |  | . | in-in |
| e(a,b) |  |  | . |

| Component 2: (a,b,(c,d),e) | | | |
| --- | --- | --- | --- |
|  | a(c,d) | b(c,d) | e(c,d) |
| a(c,d) | . | in-in | in-in |
| b(c,d) |  | . | in-in |
| e(c,d) |  |  | . |

| Component 3: ((a,b,c,d),e) | | | | | | |
| --- | --- | --- | --- | --- | --- | --- |
|  | a(b,c) | a(c,d) | a(d,e) | a,(b,d) | a,(b,e) | a,(c,e) |
| a(b,c) | . | in-out |  | in-out | in-out | in-out |
| a(c,d) |  | . | in-out | in-out |  | in-out |
| a(d,e) |  |  | . | in-out | in-out | in-out |
| a,(b,d) |  |  |  | . | in-out |  |
| a,(b,e) |  |  |  |  | . | in-out |
| a,(c,e) |  |  |  |  |  | . |

| Tree: (((a,b),(c,d)),e) (component 1 + 2 + 3) | | | | | | | | | | |
| --- | --- | --- | --- | --- | --- | --- | --- | --- | --- | --- |
|  | c(a,b) | d(a,b) | e(a,b) | a(c,d) | b(c,d) | e(c,d) | e(a,c) | e(a,d) | e(b,c) | e(b,d) |
| c(a,b) | . | in-in | in-in | s | s |  | a |  | a |  |
| d(a,b) |  | . | in-in | s | s |  |  | a |  | a |
| e(a,b) |  |  | . |  |  |  | in-out | in-out | in-out | in-out |
| a(c,d) |  |  |  | . | in-in | in-in | a | a |  |  |
| b(c,d) |  |  |  |  | . | in-in |  |  | a | a |
| e(c,d) |  |  |  |  |  | . | in-out | in-out | in-out | in-out |
| e(a,c) |  |  |  |  |  |  | . | in-out | in-out |  |
| e(a,d) |  |  |  |  |  |  |  | . |  | in-out |
| e(b,c) |  |  |  |  |  |  |  |  | . | in-out |
| e(b,d) |  |  |  |  |  |  |  |  |  | . |
