## Supplementary material for "Information Content of Trees: Three-taxon Statements Inference Rules and Dependency": Table S2

| **tr ~ number_of_permutations * number_of_taxa * analysis** |  |  | R² | 0,7501 |  |
| --- | --- | --- | --- | --- | --- |
|  | Estimate | Std. Error | t value | Pr(>\|t\|) |  |
| (Intercept) | -9,80295833 | 0,51591471 | -19,0011221 | 1,94E-80 | *** |
| number_of_permutations | 5,35899903 | 0,17046499 | 31,4375341 | 1,92E-216 | *** |
| ntax | 8,89290943 | 0,05624147 | 158,120146 | 0 | *** |
| analysisMRP | 3,65690133 | 0,72961357 | 5,01210704 | 5,39E-07 | *** |
| analysisMW | 3,76209196 | 0,72961357 | 5,15628011 | 2,52E-07 | *** |
| analysisUW | -0,18030443 | 0,72961357 | -0,24712318 | 0,80481313 | * |
| number_of_permutations:ntax | -1,38477298 | 0,01857625 | -74,5453272 | 0 | *** |
| number_of_permutations:analysisMRP | 0,85755982 | 0,2410739 | 3,55724864 | 0,00037484 | *** |
| number_of_permutations:analysisMW | 0,33260682 | 0,2410739 | 1,37968817 | 0,1676841 | *** |
| number_of_permutations:analysisUW | 0,06579629 | 0,2410739 | 0,27292995 | 0,78490732 |  |
| ntax:analysisMRP | -0,55760913 | 0,07953745 | -7,01064889 | 2,38E-12 | *** |
| ntax:analysisMW | -0,55853738 | 0,07953745 | -7,02231944 | 2,19E-12 | *** |
| ntax:analysisUW | 0,01218894 | 0,07953745 | 0,15324777 | 0,87820303 | ** |
| number_of_permutations:ntax:analysisMRP | -0,1274297 | 0,02627079 | -4,85062294 | 1,23E-06 | *** |
| number_of_permutations:ntax:analysisMW | -0,04548805 | 0,02627079 | -1,73150648 | 0,0833629 | *** |
| number_of_permutations:ntax:analysisUW | -0,01043316 | 0,02627079 | -0,39713899 | 0,69126536 | * |
| **fr ~ number_of_permutations * number_of_taxa * analysis** |  |  | R² | 0,2699 |  |
|  | Estimate | Std. Error | t value | Pr(>\|t\|) |  |
| (Intercept) | 16,8789452 | 0,52810507 | 31,9613391 | 1,26E-223 | *** |
| number_of_permutations | 2,97733672 | 0,17442736 | 17,0692067 | 2,78E-65 | *** |
| number_of_taxa | -1,16749689 | 0,07078198 | -16,4942664 | 4,40E-61 | *** |
| analysisMRP | 1,87600677 | 0,74685335 | 2,51188103 | 0,01200968 | * |
| analysisMW | 1,43302669 | 0,74685335 | 1,91875244 | 0,05501699 | . |
| analysisUW | -0,35551891 | 0,74685335 | -0,47602238 | 0,63405891 |  |
| number_of_permutations:number_of_taxa | 0,79796499 | 0,02337852 | 34,1324 | 1,17E-254 | *** |
| number_of_permutations:analysisMRP | 0,37835849 | 0,24667754 | 1,53381815 | 0,12507586 |  |
| number_of_permutations:analysisMW | 2,9412513 | 0,24667754 | 11,923466 | 9,15E-33 | *** |
| number_of_permutations:analysisUW | 0,06171054 | 0,24667754 | 0,25016685 | 0,80245856 |  |
| number_of_taxa:analysisMRP | -0,40839458 | 0,10010084 | -4,0798317 | 4,51E-05 | *** |
| number_of_taxa:analysisMW | -0,30630131 | 0,10010084 | -3,05992744 | 0,00221418 | ** |
| number_of_taxa:analysisUW | 0,10853196 | 0,10010084 | 1,08422623 | 0,27826572 |  |
| number_of_permutations:number_of_taxa:analysisMRP | -0,10271017 | 0,03306222 | -3,10657213 | 0,00189295 | ** |
| number_of_permutations:number_of_taxa:analysisMW | -0,56976281 | 0,03306222 | -17,2330477 | 1,66E-66 | *** |
| number_of_permutations:number_of_taxa:analysisUW | -0,01064829 | 0,03306222 | -0,32206807 | 0,74740144 |  |
| **efficiency ~ number_of_permutations * number_of_taxa * analysis** | |  | R² | 0,5712 |  |
|  | Estimate | Std. Error | t value | Pr(>\|t\|) |  |
| (Intercept) | 18,1816187 | 0,32726977 | 55,5554478 | 0 | *** |
| number_of_permutations | 0,63479849 | 0,10809365 | 5,87267155 | 4,29E-09 | *** |
| number_of_taxa | 7,35606337 | 0,04386401 | 167,701589 | 0 | *** |
| analysisMRP | 2,31814092 | 0,46282935 | 5,00862992 | 5,49E-07 | *** |
| analysisMW | 3,45524832 | 0,46282935 | 7,46549096 | 8,33E-14 | *** |
| analysisUW | -0,32082461 | 0,46282935 | -0,69318122 | 0,48819658 |  |
| number_of_permutations:number_of_taxa | -1,0924965 | 0,0144878 | -75,4080119 | 0 | *** |
| number_of_permutations:analysisMRP | 0,58142376 | 0,1528675 | 3,80344909 | 0,00014273 | *** |
| number_of_permutations:analysisMW | 0,16060499 | 0,1528675 | 1,05061563 | 0,29343633 |  |
| number_of_permutations:analysisUW | 0,08873633 | 0,1528675 | 0,5804787 | 0,56159245 |  |
| number_of_taxa:analysisMRP | -0,52038693 | 0,06203307 | -8,38886298 | 4,94E-17 | *** |
| number_of_taxa:analysisMW | -0,6690952 | 0,06203307 | -10,7861047 | 4,07E-27 | *** |
| number_of_taxa:analysisUW | 0,03564742 | 0,06203307 | 0,57465193 | 0,56552732 |  |
| number_of_permutations:number_of_taxa:analysisMRP | -0,12356586 | 0,02048885 | -6,03088344 | 1,63E-09 | *** |
| number_of_permutations:number_of_taxa:analysisMW | -0,03371683 | 0,02048885 | -1,64561866 | 0,09984375 | . |
| number_of_permutations:number_of_taxa:analysisUW | -0,01660792 | 0,02048885 | -0,81058352 | 0,41760578 |  |
