## Supplementary figures and images for "Information Content of Trees: Three-taxon Statements Inference Rules and Dependency"

### Figure S1

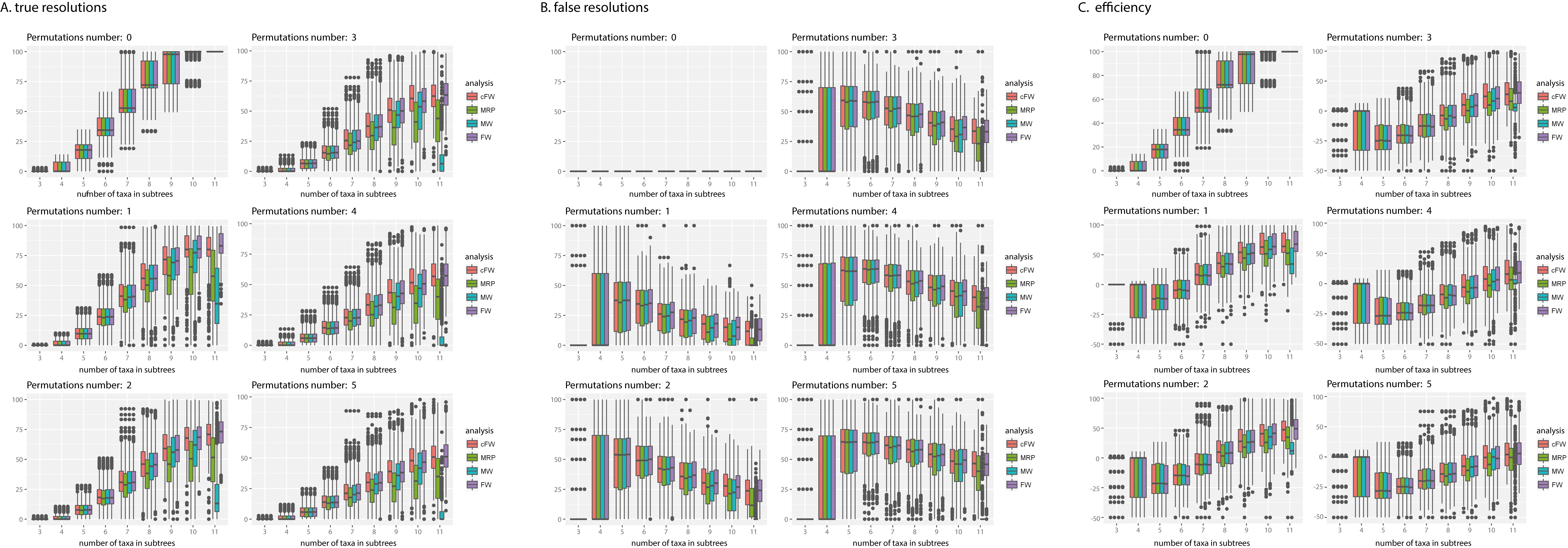

### Figure S3

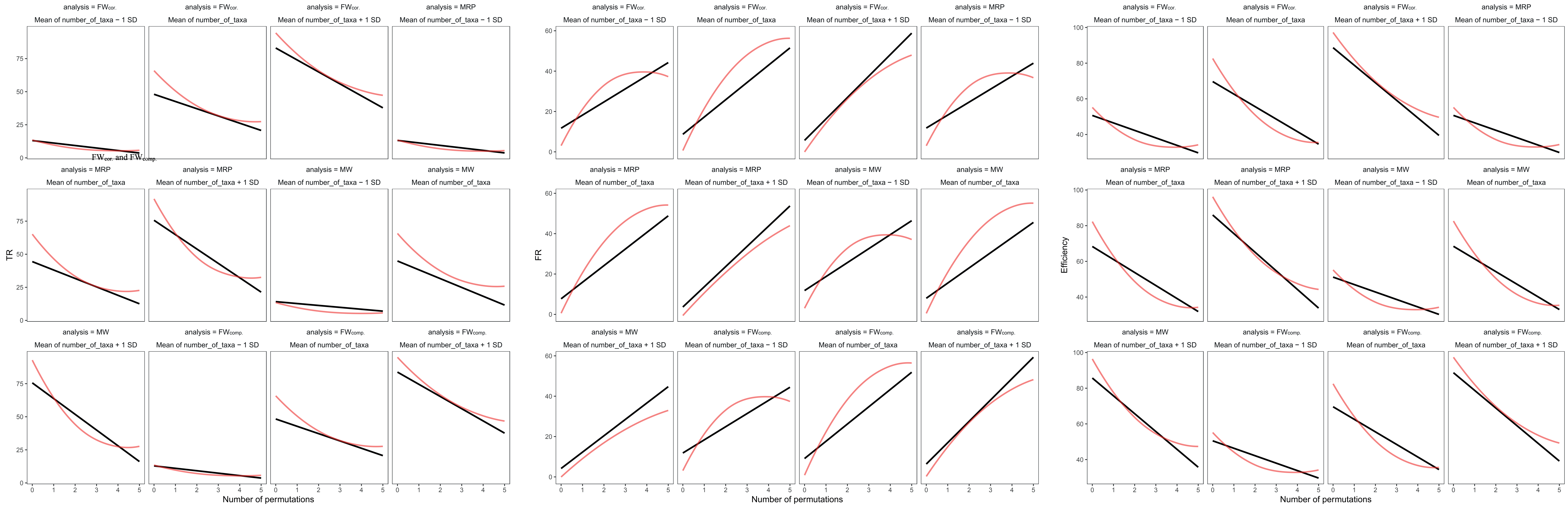
